# The role of Cycloastragenol at the intersection of Nrf-2/ARE, telomerase, and proteasome activity

**DOI:** 10.1101/2022.03.11.483898

**Authors:** Sinem Yılmaz, Erdal Bedir, Petek Ballar Kırmızıbayrak

## Abstract

Aging is well-characterized by the gradual decline of cellular functionality. As redox balance, proteostasis, and telomerase systems have been found to be associated with aging and age-related diseases, targeting these systems by small compounds has been considered as a promising therapeutic approach. Cycloastragenol (CA), a small molecule telomerase activator obtained from *Astragalus* species, has been reported to have positive effects on several age-related pathophysiologies, including ischemia, glucose intolerance, diabetes, and neurodegenerative diseases. Although CA has received intense attention as a promising compound for aging and age-related diseases, the mechanisms underlying CA activity related to redox balance, proteasome, and telomerase has not yet been reported. Here, we presented that CA increased Nrf-2 activity leading to upregulation of cytoprotective enzymes and attenuation of oxidative stress-induced ROS levels. Furthermore, CA enhanced telomerase activity by increasing not only the expression of hTERT but also its nuclear localization via the Hsp90-chaperon complex. CA alleviated the oxidative stress-induced mitochondrial hTERT levels while increasing the nuclear hTERT levels. Additionally, the proteasome activity and assembly were increased by CA. Strikingly, our data revealed that CA-mediated neuroprotection requires both Nrf-2 and hTERT induction. In conclusion, this study is the first report describing the effect of CA on these aging-related three major cellular pathways and their interrelationships. As the proteasome activator effect of CA is dependent on induction of telomerase activity, which is mediated by Nrf-2 system, CA has a great potential for healthier aging and prevention or treatment of age-related diseases by positively affecting these three cellular pathways.

## 1. INTRODUCTION

Aging is associated with a general decrease in cellular functionality. Several hallmarks such as genomic instability, telomere attrition, epigenetic alterations, decline of proteolysis, mitochondrial dysfunction, cellular senescence, stem cell exhaustion, and altered intercellular communication have been described for aging cells (Lopez-Otin et al., 2013). These changes not only lead to acceleration of the progression of aging but also to the development of age-related pathologies such as cardiovascular diseases, diabetes, and neurodegenerative diseases. Telomere shortening, increased reactive oxygen species (ROS) formation, and decline of proteasomal activity are three main conserved cellular alterations upon aging that have been focus of research to promote healthier aging (Lopez-Otin et al., 2013).

The accumulation of oxidative damage is one of the critical factors during aging, and its alleviation has been shown to extend the lifespan (Salmon et al., 2010). The Nrf-2 (nuclear factor erythroid 2–related factor 2)/Keap-1 (Kelch-like ECH-associated protein 1) signaling pathway is a major cellular defense mechanism against oxidative stress (Salmon et al., 2010). While Nrf-2 is a redox sensing transcription factor and coordinates many cytoprotective genes, mainly including Antioxidant Response Elements (AREs) promoter region (Schmidlin et al., 2019; Zhang, H.Q. et al., 2015), Keap-1 is the negative regulator of this signaling pathway, and its function depends on the intracellular redox state (Zhang, H. et al., 2015). Under unstressed conditions, Keap-1 sequesters Nrf-2 in the cytoplasm and promotes its ubiquitination and proteasomal degradation. Increased oxidative stress induces a conformational change in the Nrf-2-Keap-1 complex preventing the ubiquitination of Nrf-2 and promoting its nuclear translocation (Zhang, D.D. & Hannink, 2003). It is well known that Nrf-2 inactivation exacerbates the progression of several aging-related phenotypes and dysfunctions (Ahn et al., 2018; Fulop et al., 2018; Gounder et al., 2012; Hecker et al., 2014; Narasimhan et al., 2014; Tarantini et al., 2018; Valcarcel-Ares et al., 2017; Yoh et al., 2001). As several studies describe that Nrf-2 activity is decreased upon aging, and enhanced Nrf-2 activity correlates with extended lifespan in various organisms, Nrf-2 has been implicated as a therapeutic target in age-related diseases (Chapple et al., 2012).

The ability of the cell to maintain its protein homeostasis (proteostasis) is also affected upon aging progression. The proteasome, a multi-catalytic enzyme complex located in the cytosol and nucleus, is formed by the assembly of the barrel-shaped proteolytic core particle (20S) and the regulatory particle (19S) (Saez & Vilchez, 2014). Proteasome, which is considered as the main component for surveilling proteostasis, is responsible for degradation of misfolded, unfolded, damaged, short-lived, or regulatory proteins, which are modified by a poly-ubiquitin chain that facilitates their recognition by the proteasome. In addition to ubiquitin-dependent proteasomal degradation, a certain fraction of unfolded and intrinsically disordered proteins are also degraded by proteasome in a ubiquitin-independent manner, especially under oxidative stress (Baugh et al., 2009; Ben-Nissan & Sharon, 2014). The activity of the proteasome decreases dramatically with aging (Andersson et al., 2013; Chondrogianni et al., 2015). The accompanied progressive accumulation of damaged or misfolded proteins by proteasomal dysfunction is increased to lead to aging and/or age-related diseases. Hence, the inhibition of proteasome in young cells induced premature senescence-like phenotype (Bellavista et al., 2014). Downregulation of proteosome subunit expressions, alterations in post-translational modification of subunits, disruption of proteasome assembly, and occlusion of the proteasome by aggregated or cross-linked proteins are reported factors responsible for the reduction of proteosome activity observed during the aging process (Chondrogianni et al., 2014; Vilchez et al., 2014). Thus, several approaches to enhance proteasome activity such as genetic manipulations, post-translational modifications and small molecule proteasome agonists have been depicted to prevent or decelerate the progression of age-related diseases such as Parkinson’s and Alzheimer’s and extend health span and longevity (Chondrogianni et al., 2014; Chondrogianni et al., 2005; Chondrogianni et al., 2015; Kapeta et al., 2010).

Telomeres, consisting of TTAGGG repeats, are located at the ends of linear chromosomes and become progressively shorter upon each cell division (Greider, 1996). Telomerase, which contains a telomerase reverse transcriptase (TERT) and the Telomerase RNA Component (TERC), reverse transcribes telomeres and retards cellular aging (Blackburn, 1991). TERT expression and telomerase activity are very low or absent in most multicellular eukaryotic organisms but are significantly high in stem cells and germ cells. Furthermore, recently it has been depicted that TERT is also present in tissues with low replicative potential, such as the heart (Richardson et al., 2012; Zurek et al., 2016) and the brain (Iannilli et al., 2013; Spilsbury et al., 2015). Continuingly, non-canonical functions of TERT have been very recently described, including signal transduction, gene expression regulation, protection against oxidative damage, and chaperone activity for the 26S proteasome assembly that are independent of its telomere elongation activity (Segal-Bendirdjian & Geli, 2019). Considering its telomere-dependent and independent roles of TERT, its activation with potent compounds is important for developing new strategies in age-related diseases.

Cycloastragenol (CA, 20(R),24(S)-epoxy-3β,6α,16β,25-tetrahydroxycycloartane) is a small molecule telomerase activator derived from the extract of *Astragalus membranaceous*, a plant commonly used in traditional Chinese medicine to treat several diseases such as diabetes, hyperlipidemia, atherosclerosis, and cancer. CA not only increases telomerase activity and elongates telomere length but also extends life span of mice without augmenting cancer incidences (Bernardes de Jesus et al., 2011). Thus, CA and related derivatives have great potential as anti-aging agents and treatment of age-related diseases. Indeed, CA (or CA-containing TA-65®) has been shown to have beneficial effects on several conditions such as glucose intolerance, insulin resistance, immunity, neural depression, age-related macular degeneration, and cardiovascular diseases (Dow & Harley, 2016; Harley et al., 2013; Hummasti & Hotamisligil, 2010; Ip et al., 2014; Libby et al., 2010; Zhao et al., 2015). Furthermore, several clinical studies suggest that, as a new dietary supplement, TA-65^®^ may enhance one’s health span without any adverse effects (Harley et al., 2013). Despite all the promising effects of CA, which were demonstrated by several studies, the underlying molecular mechanisms of how CA affected telomerase and redox balance were missing. In this study, we report for the first time that proteasome activator action of CA is dependent on induction of telomerase activity, which is driven by activation of Nrf-2 system. Therefore, our results revealed that CA treatment affects multiple interconnected aging-related cellular processes such as hTERT, Nrf-2, and proteasome, suggesting that CA is a highly prominent candidate for the treatment and/or prevention of aging and aging-related diseases.

## 2. RESULTS

### 2.1. CA is a novel Nrf-2 activator

The effect of CA on Nrf-2 signaling was first investigated via nuclear transcriptional activity of Nrf-2 in young HEKn (passage 5). The treatment of CA increased both protein and mRNA levels of Nrf-2 in a dose-dependent manner (**Figure 1a,b**). Furthermore, immunofluorescence and cellular fractionation assays revealed that CA promoted nuclear localization of Nrf-2 from cytoplasm (**Figure 1c,d**). These observations are well-consistent with our result showing a dose-dependent augmentation of the nuclear Nrf-2 activity starting as low as 0.5 nM (**Figure 1e**). The effect of CA on the Nrf-2 activity was much more prominent in older HEKn cells at passage 15, which had entered senescence as evidenced by the expression of senescence-related p16 (**Figure S1A,B**). As our results suggest that CA might be a novel Nrf-2 activator, next ARE promoter activity was analyzed. Even at nanomolar concentrations, CA enhanced the relative luciferase activity more than DMF (50 μM, Dimethyl fumarate), a well-known Nrf-2 activator (**Figure 1f**). While CA enhanced the protein level of HO-1 (Heme oxygenase 1), one of the downstream components of Nrf-2 signaling, it did not significantly alter protein levels of a negative regulator, Keap-1 (**Figure 1g**). We further investigated the effect of CA on the activity of the cytoprotective enzymes and found that CA increased the activities of HO-1, GR (Glutathione reductase), and GCLC (Glutamate-cysteine ligase catalytic subunit) enzymes (**Figure 1h**). Consistent with these results, CA treatment dose-dependently alleviated H_2_O_2_-induced ROS formation (**Figure 1i**). Taking together, our results point out that CA treatment promotes Nrf-2 activation, and this activity leads to upregulation of several cytoprotective enzymes that protect cells against oxidative stress damage.

**FIGURE 1.**
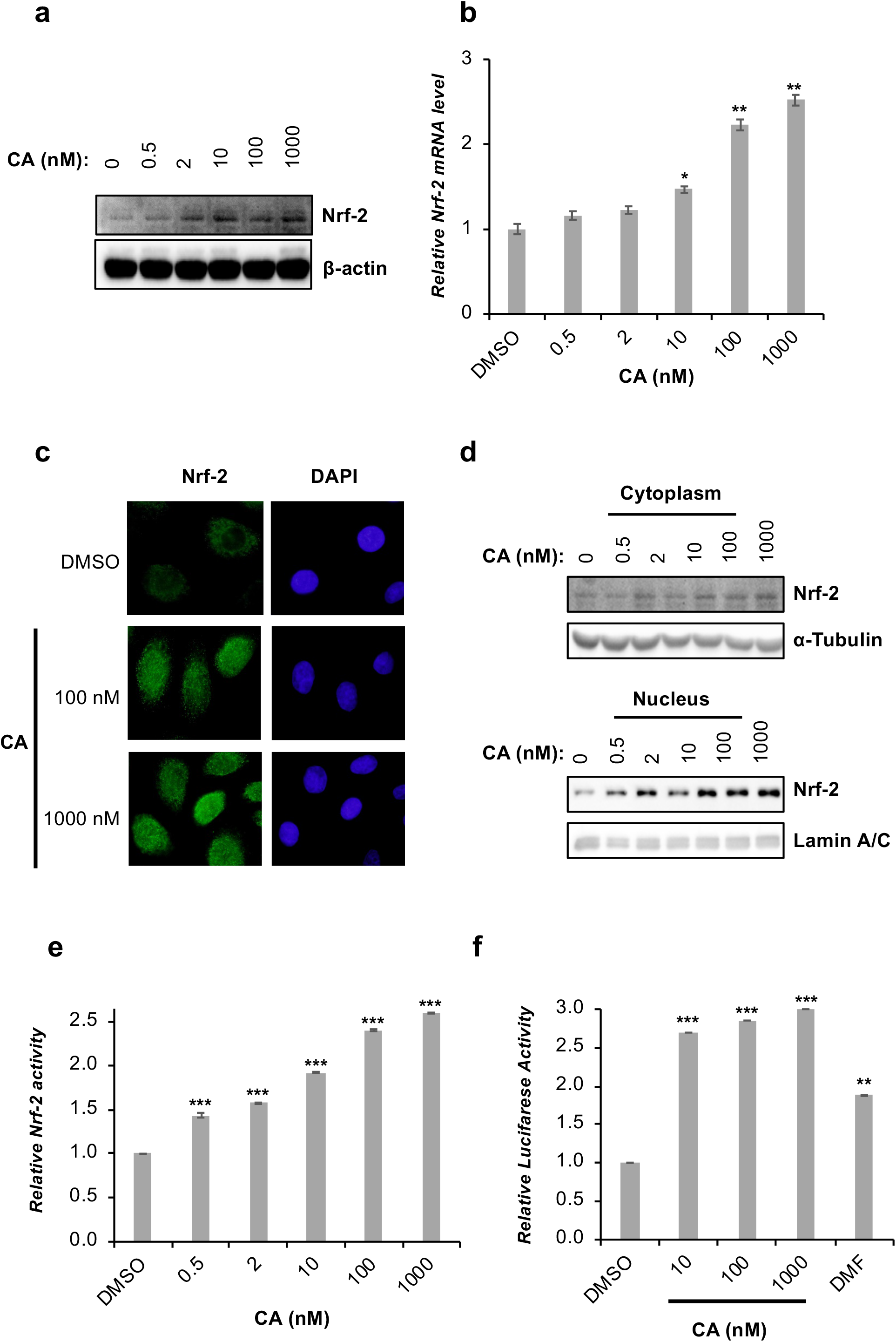
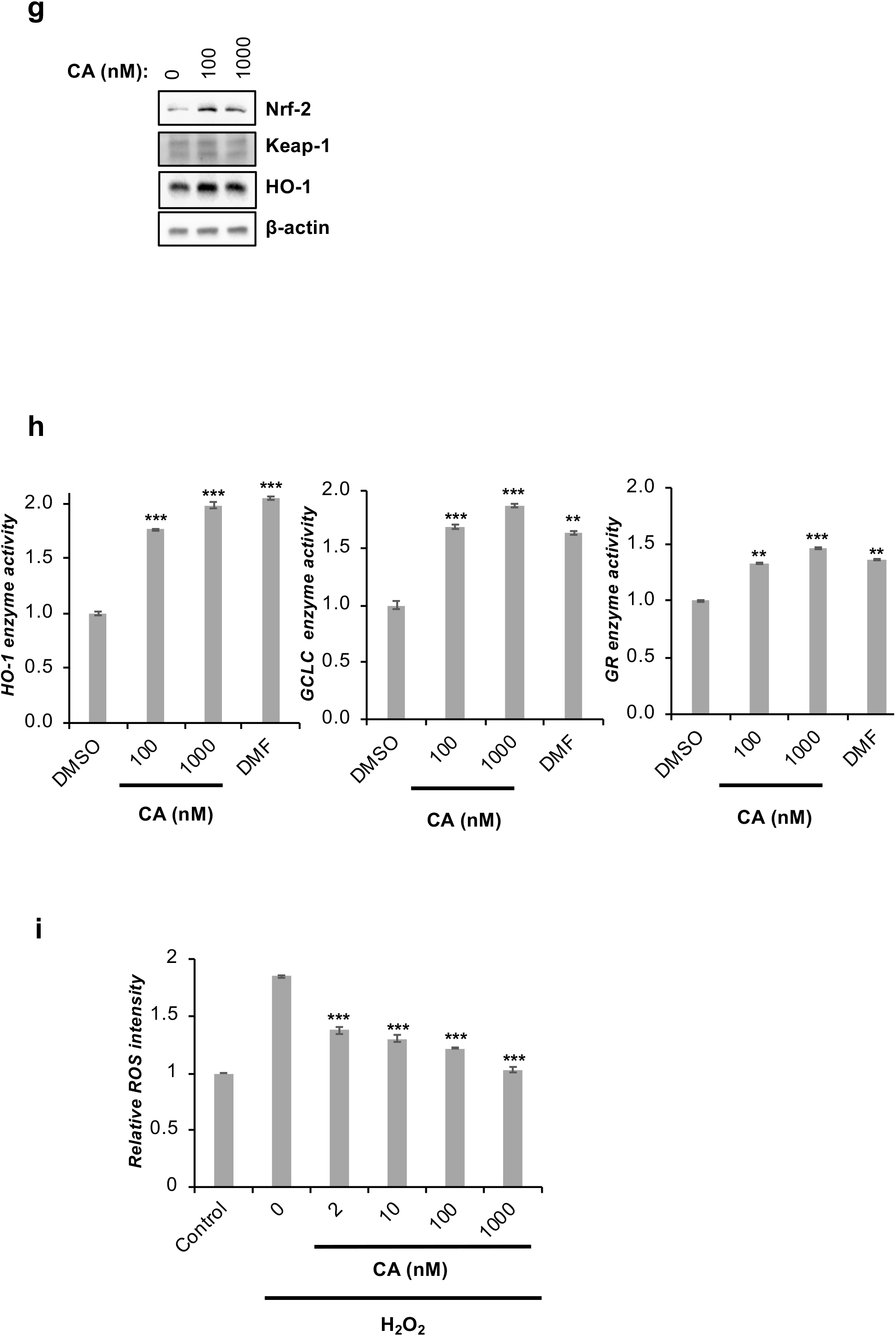
The effect of CA on Nrf-2/ARE system. HEKn cells were treated with indicated concentrations of CA for 24 h. (**a**) Nrf-2 protein levels were determined by IB. β-actin was used as a loading control. (**b**) The mRNA levels of Nrf-2 were investigated by RT-qPCR. Error bars are presented standard error (n=3, *p ≤ 0.05, **p ≤ 0.001, ***p ≤ 0.005). (**c**) The subcellular localization of Nrf-2 was evaluated by immunostaining using anti-Nrf-2 antibody. Representative fluorescent images for nuclear translocation of Nrf2. DAPI was used for nuclei staining. (**d)** Cellular fractionation assay was performed, and Nrf-2 protein levels were investigated by IB. While Lamin A-C was used as nuclear loading control, α-Tubulin was used as cytoplasmic loading control. (**e**) Nrf-2 transcription factor assay using the nuclear proteins obtained from control or CA-treated cells was carried out by an ELISA kit. The obtained nuclear activity data was normalized to nuclear protein levels and presented as a fold change compared to control cells treated with DMSO. Error bars are presented as standard deviations (n = 3; *p ≤ 0.05, **p ≤ 0.001, ***p ≤ 0.005). (**f**) The ARE-promotor activity was determined by Luciferase activity assay. DMF (50 μM) was used as an experimental control. Error bars are presented as standard deviations (n = 3; *p ≤ 0.05, **p ≤ 0.001, ***p ≤ 0.005). (**g**) HO-1, Keap-1, and Nrf-2 protein levels were investigated by IB. β-actin was used as a loading control. (**h**) HO-1, GCLC and GR cytoprotective enzymes levels were quantified by ELISA. DMF (50 μM) was used as an experimental control. The enzyme activity levels were normalized to cellular total protein levels and presented as a fold change compared to control cells treated with DMSO. Error bars are presented as standard deviations (n = 3; *p ≤ 0.05, **p ≤ 0.001, ***p ≤ 0.005). (**i**) The effect of ROS (induced by H_2_O_2_) scavenging properties of CA was determined by H_2_DCFDA. Error bars are presented as standard deviations (n = 3; *p ≤ 0.05, **p ≤ 0.001, ***p ≤ 0.005).

### 2.2. CA activates telomerase by enhancing not only hTERT expression but also its nuclear localization

Several studies have suggested that TERT mRNA expression associated with telomerase activation is elevated in response to CA treatment in primary cortical neurons, human epidermal stem cells, osteoblastic MC3T3-E1 cells, and *in vivo* in some mouse tissues (Bernardes de Jesus et al., 2011; Cao et al., 2019; Ip et al., 2014; Wu et al., 2021). Consistently, CA treatment promoted both hTERT protein and mRNA levels in a dosedependent manner in young HEKn cells (**Figure 2a,b**). CA-mediated elevation of hTERT levels was also prominent in senescent HEKn cells (**Figure S2**). As TERT must translocate to the nucleus to form catalytically active telomerase (Jeong, Y.Y. et al., 2016), we next evaluated the localization of hTERT upon CA treatment and have found that CA enhanced hTERT translocation to the nucleus (**Figure 2c**). After this observation, we further evaluated the precise molecular mechanism underlying CA-induced nuclear transport of hTERT by analyzing the effect of CA on the levels of several proteins that act as regulators of hTERT nuclear translocation. The level of CHIP (C terminus of Hsc70-interacting protein), which prevents the nuclear localization of hTERT and facilitates its degradation (Lee et al., 2010), was decreased in CA-treated HEKn cells (**Figure 2d, left, lane 2 vs. 1**). The decrease in CHIP expression was also observed in senescent HEKn cells (**Figure S3A**). On the other hand, the levels of regulatory proteins Hsp90 (Heat shock protein 90), p23, and FKBP52 (FK506-binding protein 52), which function in the hTERT nuclear translocation complex, were enhanced in response to CA treatment (**Figure 2d, right, lane 2 vs. 1**). Moreover, the increase in the levels of Hsp90, p23, hTERT, and FKBP52 was also observed in MRC-5 cell line (normal lung fibroblast cell line) (**Figure S3B**). As geldanamycin (GA), a well-known Hsp90 inhibitor, was reported to inhibit telomerase activity through the ubiquitin-mediated degradation of hTERT (Kim et al., 2005), we next pre-treated cells with GA prior to CA treatment. Consistently, our results revealed that GA ameliorated the CA-mediated upregulation of Hsp90, p23, and FKBP52 (**Figure 2d, lane 3 vs. 2**). Furthermore, GA inhibited CA-mediated nuclear translocation of hTERT as well as Hsp90-based heterocomplex (**Figure 2e**). As the interaction of Hsp90 and hTERT is essential both for proper assembly of active telomerase and for nuclear import of hTERT (Forsythe et al., 2001; Holt et al., 1999; Jeong, S.A. et al., 2015), we next evaluated the effect of CA on the Hsp90-hTERT interaction and found that CA significantly enhanced hTERT-Hsp90 interaction (**Figure 2f**).

**FIGURE 2.**
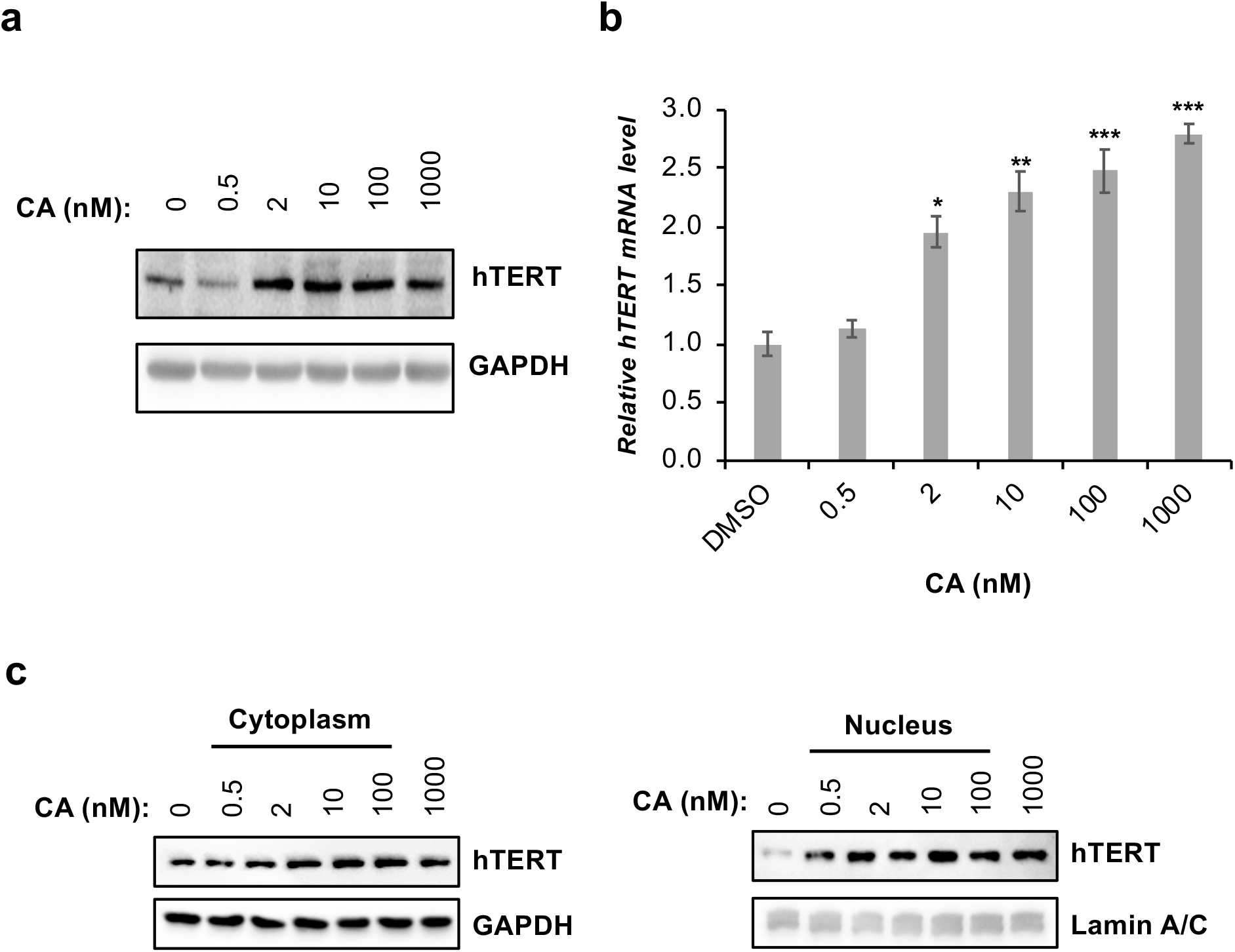
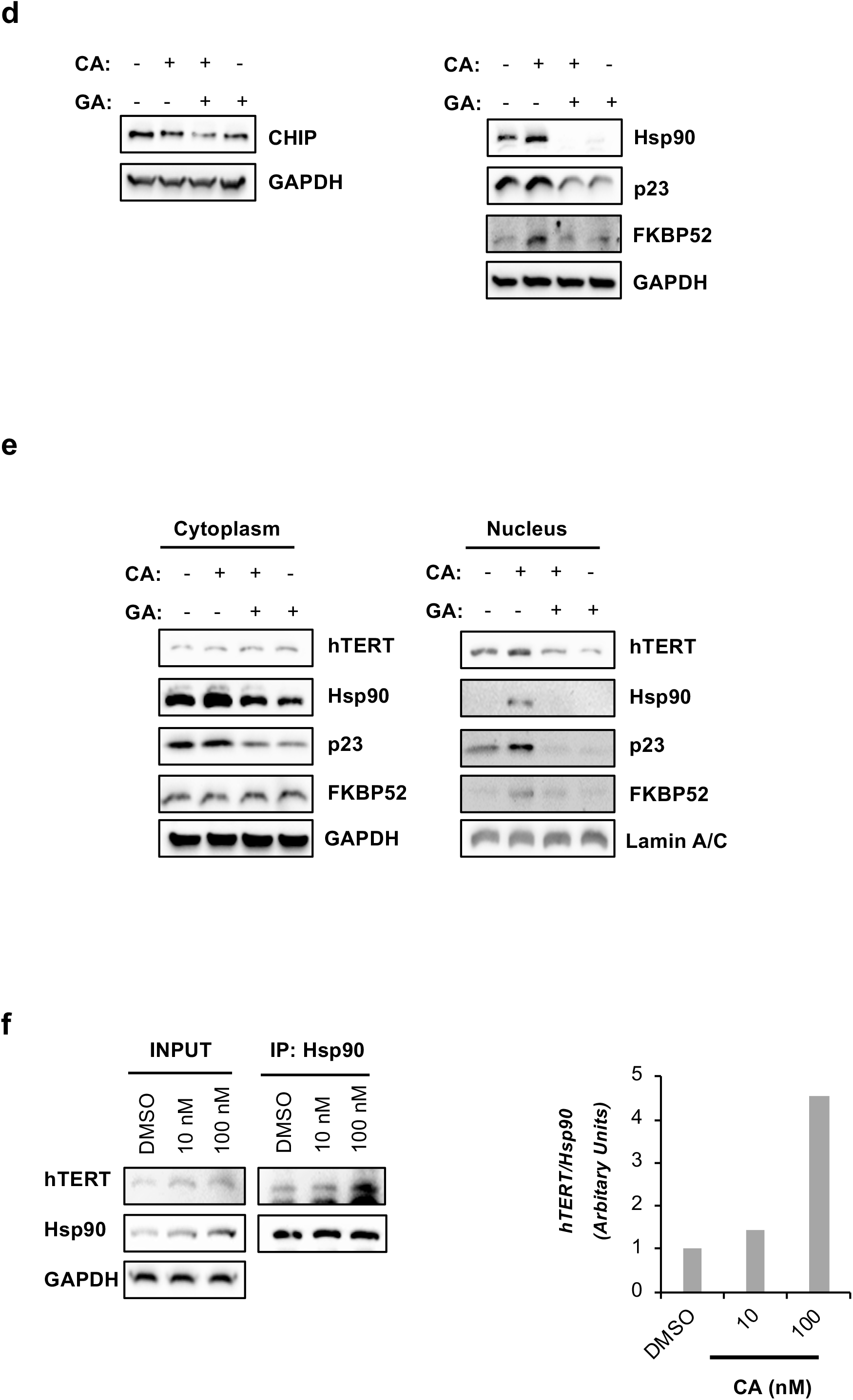
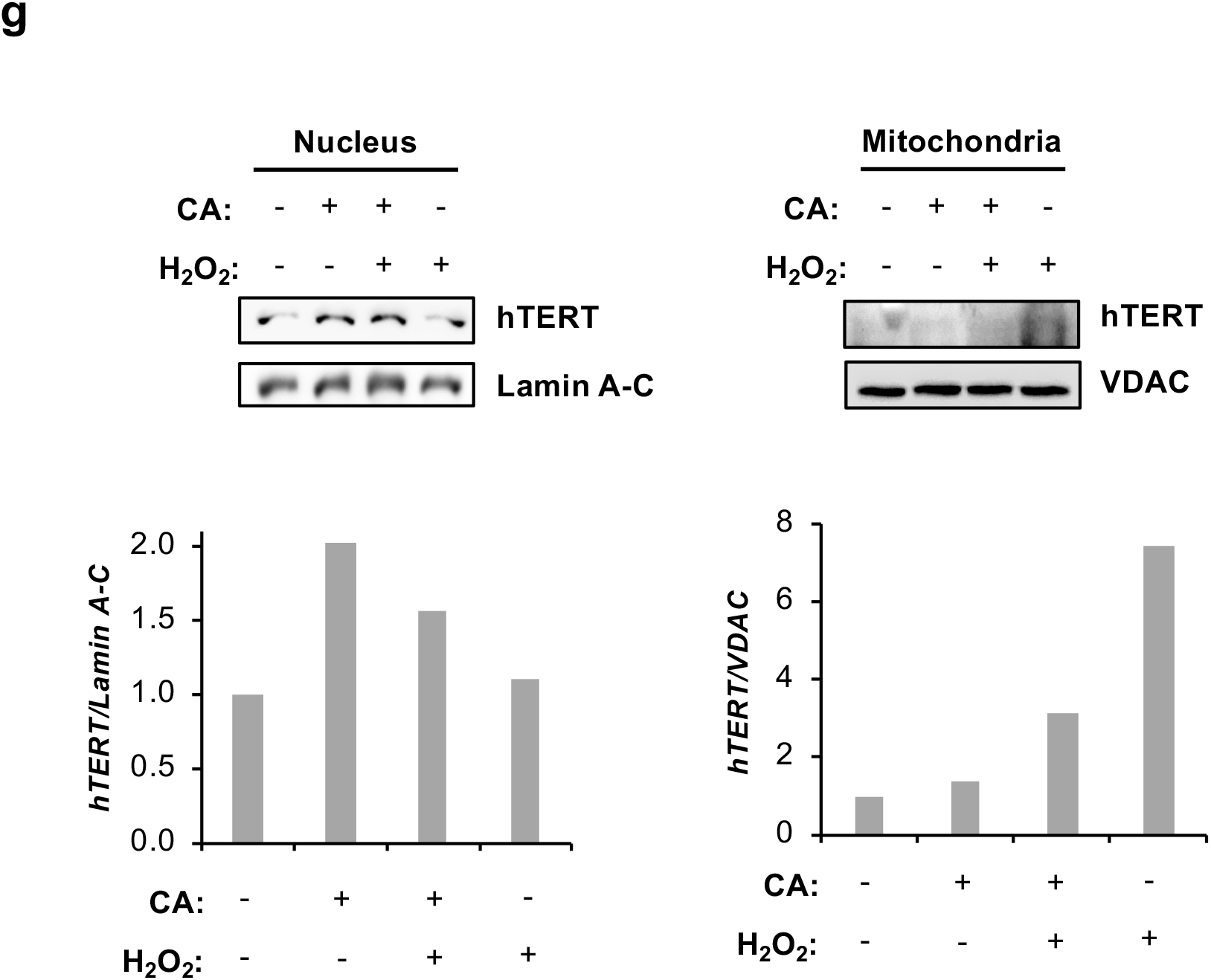
CA increased not only mRNA and protein levels of hTERT but also nuclear localization of hTERT *via* Hsp90-chaperon complex. **(a, b, c)** HEKn cells were treated with indicated concentrations of CA for 24 h treatment. (**a**) The protein levels of hTERT were determined by IB. GAPDH was used as a loading control. (**b**) The mRNA levels of hTERT were investigated by RT-qPCR. Error bars are presented as standard deviations (n = 3; *p ≤ 0.05, **p ≤ 0.001, ***p ≤ 0.005). (c) The effect of CA on the hTERT nuclear localization was determined by cellular fractionation assay. hTERT protein levels were determined by IB. While Lamin A-C was used as nuclear loading control, α-Tubulin was used as cytoplasmic loading control. (**d, e, f**) HEKn cells were pre-treated with 100 nM GA known as Hsp90 inhibitor for 1 h, then treated with CA for an additional 24 h. (**d**) Total protein levels of CHIP, Hsp90, p23, and FKBP52 were determined by IB. (**e**) The cellular fractionation was performed as in cytoplasmic and nuclear protein levels of hTERT, Hsp90, p23, and FKBP52 were determined by IB. (**f**) The effect of CA on Hsp90-hTERT association was analyzed by immunoprecipitation. The protein levels of Hsp90 and hTERT were evaluated by IB. GAPDH was used as a loading control for Input. (**g**) HEKn cells were pre-treated with CA for 1 h then treated with H_2_O_2_ for an additional 24 h. Nuclear and mitochondrial fractionation was performed. Mitochondrial and nuclear hTERT levels were determined by IB. While Lamin A-C was used as nuclear loading control, VDAC was used as a mitochondrial loading control.

Over the last years, non-canonical functions of hTERT have been reported, including controlling ROS levels in the cytosol and mitochondria, which is suggested to be closely related to the presence of hTERT also in mitochondria (Rosen et al., 2020). As hTERT is increased in the mitochondria upon short-term oxidative stress (Haendeler et al., 2003; Rosen et al., 2020), the effect of CA on the mitochondrial hTERT levels was investigated. As previously reported, H_2_O_2_ induced the mitochondrial but not nuclear hTERT levels (**Figure 2g**, **lane 1 vs. 4**). Our data also revealed that CA treatment increased the levels of nuclear hTERT but showed no significant change in mitochondrial hTERT levels in HEKn cells under basal conditions (**Figure 2g**, **lane 2 vs. 1**). Furthermore, CA alleviated the H_2_O_2_-induced mitochondrial hTERT levels while increasing the nuclear hTERT levels decreased by H_2_O_2_ in HEKn cells. In conclusion, our results strongly suggested that CA increased not only the hTERT expression but also the levels of components of Hsp90-based heterocomplex and enhanced the nuclear localization of hTERT.

### 2.3. CA enhances proteasomal activity

Based on prior findings that both Nrf-2 and hTERT are associated with the regulation of proteasomal activity (Imai et al., 2003; Kapeta et al., 2010), we hypothesized that CA might activate proteasome as it enhanced Nrf-2 activity and hTERT levels. To verify this hypothesis, we next evaluated the effect of CA on the proteasomal activity by using fluorogenic substrates specific to each proteasome subunit. The incubation of different concentrations of CA with the proteasome-enriched lysates of HEKn did not affect the chymotrypsin-(β5), trypsin-(β2), and caspase-like (β1) activities (**Figure 3a**), while proteasome inhibitor Mg-132 decreased the activity of each proteasomal subunit as expected (**Figure S4**). These observations suggested that CA did not directly target proteasomal activity. Next, HEKn cells were treated with CA for 24 hours, and then the status of proteasomal activity was evaluated. While CA did not significantly affect trypsin-like activity, it significantly increased both chymotrypsin- and caspase-like activities in a dose-dependent manner (**Figure 3b**). Furthermore, CA treatment increased the expression levels of β1 and β5 genes, but not β2, in a dose-dependent manner (**Figure 3c, d**). Importantly, the effect of CA to increase proteasomal subunit expressions and proteasomal activity was also observed in senescent HEKn cells (**Figure S5A,B**). The analysis of proteasome assembly by proteasome immunoprecipitation revealed that the assembly of proteasome was enhanced following CA treatment, further demonstrating that CA induces proteasome activity (**Figure 3e**). Importantly, 18α-GA (18α-Glycyrrhetinic acid), reported as a proteasome activator, enhanced proteasome assembly at micromolar concentrations (Kapeta et al., 2010), while CA at lower nanomolar concentrations suggesting that CA might have a better therapeutic index than 18α-GA.

**FIGURE 3.**
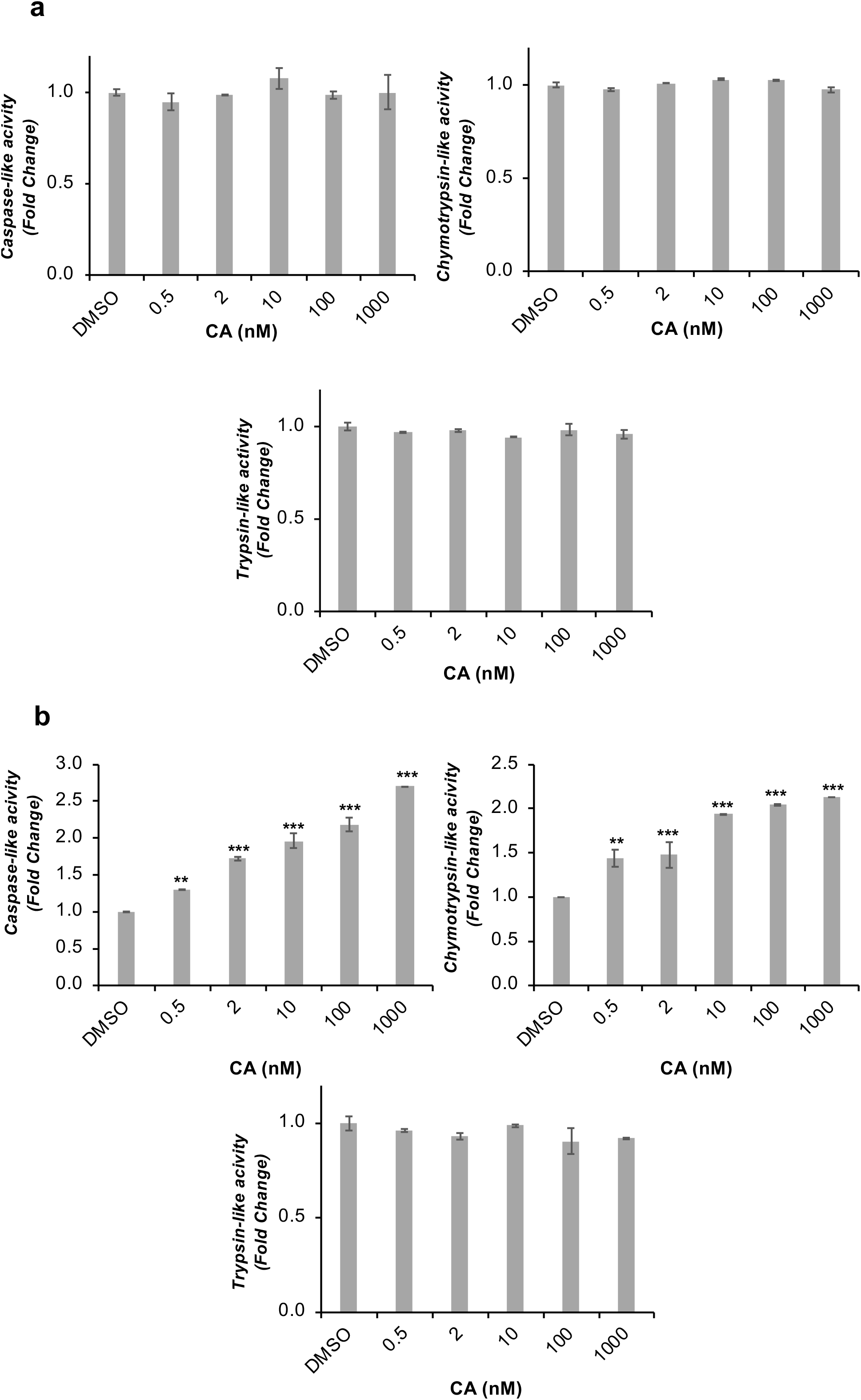
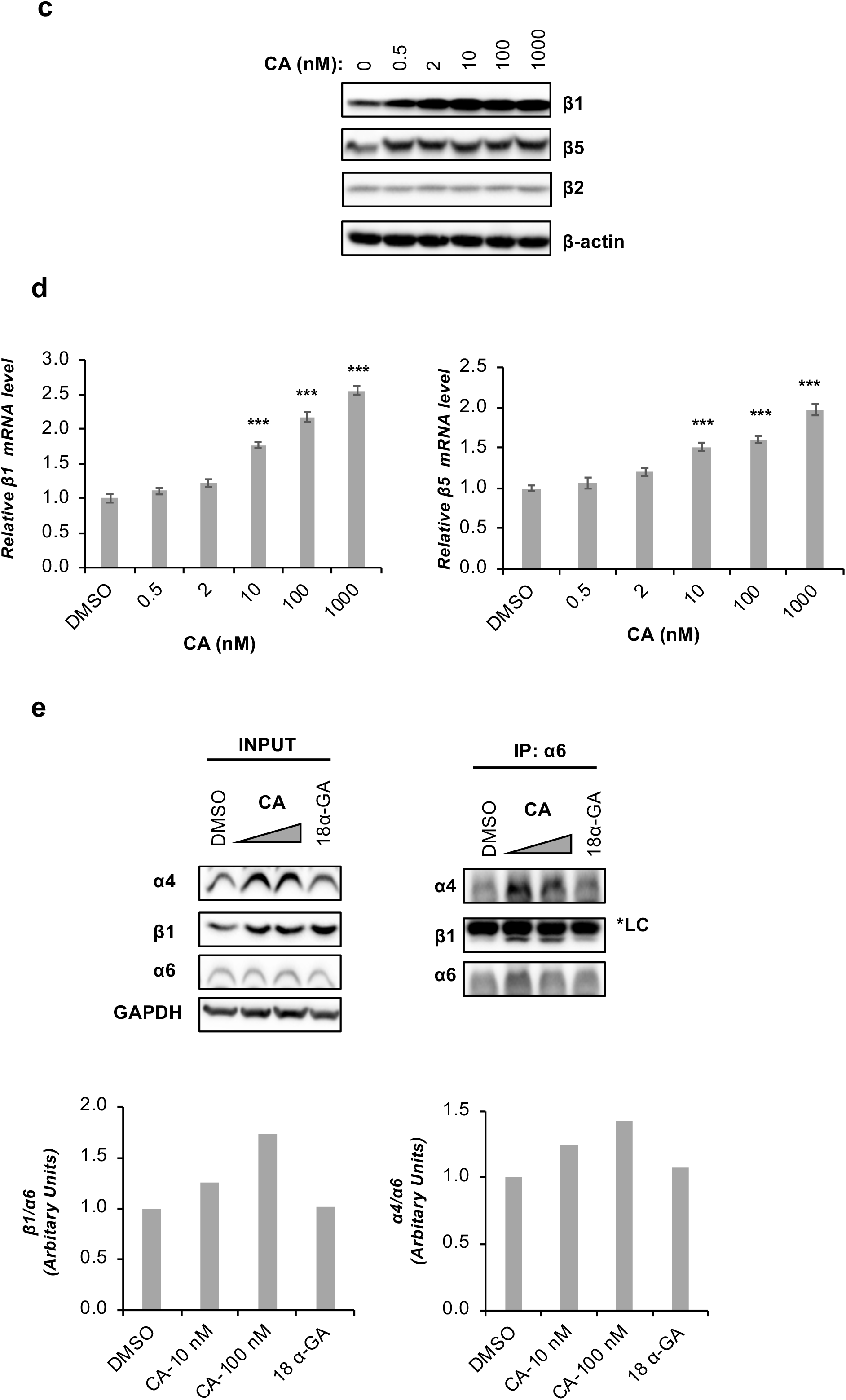
The effect of CA on proteasome status. **(a)** Proteasome-enriched lysate of HEKn cells was incubated with indicated concentrations of CA. Proteasome subunits’ activities were determined *via* fluorogenic substrates (n = 3; *p ≤ 0.05, **p ≤ 0.001, ***p ≤ 0.005). (**b, c, d, e**). HEKn cells were treated with the indicated concentration of CA for 24 h. (**b**) Fluorogenic substrates were used to evaluate the proteasome subunits’ activities. The activity data were normalized to cellular total protein level and presented as a fold change compared to control cells treated with DMSO used as solvent control. Error bars are presented as standard deviations (n = 3; *p ≤ 0.05, **p ≤ 0.001, ***p ≤ 0.005). (**c**) The total protein levels of β1, β2, and β5 were evaluated by IB. β-actin was used as a loading control. (**d**) The mRNA levels of β1 and β5 were investigated by RT-qPCR. Error bars are presented as standard deviations (n = 3; *p ≤ 0.05, **p ≤ 0.001, ***p ≤ 0.005). (**e**) The effect of CA on the proteasome assembly was determined by IP. 18 α-GA (2 μg/ml; 4.25 μM) was used as an experimental control. After IP, the protein levels of α6, α4, and β1 were evaluated by IB using antibodies against them; *LC, light chain. GAPDH was used as a loading control for Input.

### 2.4. Nrf-2 and telomerase mediated-proteasome activation by CA

Considering that CA increased the nuclear transport and activity of Nrf-2 and hTERT, which are associated with proteasomal activity, we next focused on the possible interrelationship of these three-biological processes in the context of CA. When Nrf-2 expression was silenced, the expression levels of hTERT and β1 and β5 proteasomal subunits were decreased (**Figure 4a**). Interestingly, hTERT silencing had no significant effect on Nrf-2 levels but abolished β1 and β5 protein expression (**Figure 4a**). Additionally, CA was no longer able to enhance hTERT, β1, and β5 protein levels in Nrf-2 silenced HEKn cells (**Figure 4b**). Furthermore, Nrf-2 silencing completely attenuated CA-induced telomerase and proteasome activities (**Figure 4c,d**). These data strongly suggest that the CA-mediated increase in telomerase and proteasome activities is driven by Nrf-2 activity. Interestingly, we observed that CA enhanced Nrf-2 protein levels but not proteasome subunit levels in hTERT silenced HEKn cells (**Figure 5a**). Consistently, CA was able to enhance Nrf-2 nuclear activity, but not proteasome activity in hTERT silenced cells (**Figure 5b,c**). These data suggest that the proteasome activator ability of CA is mediated by hTERT, and the enhancement of telomerase activity and hTERT levels by CA treatment is driven by Nrf-2 system.

**FIGURE 4.**
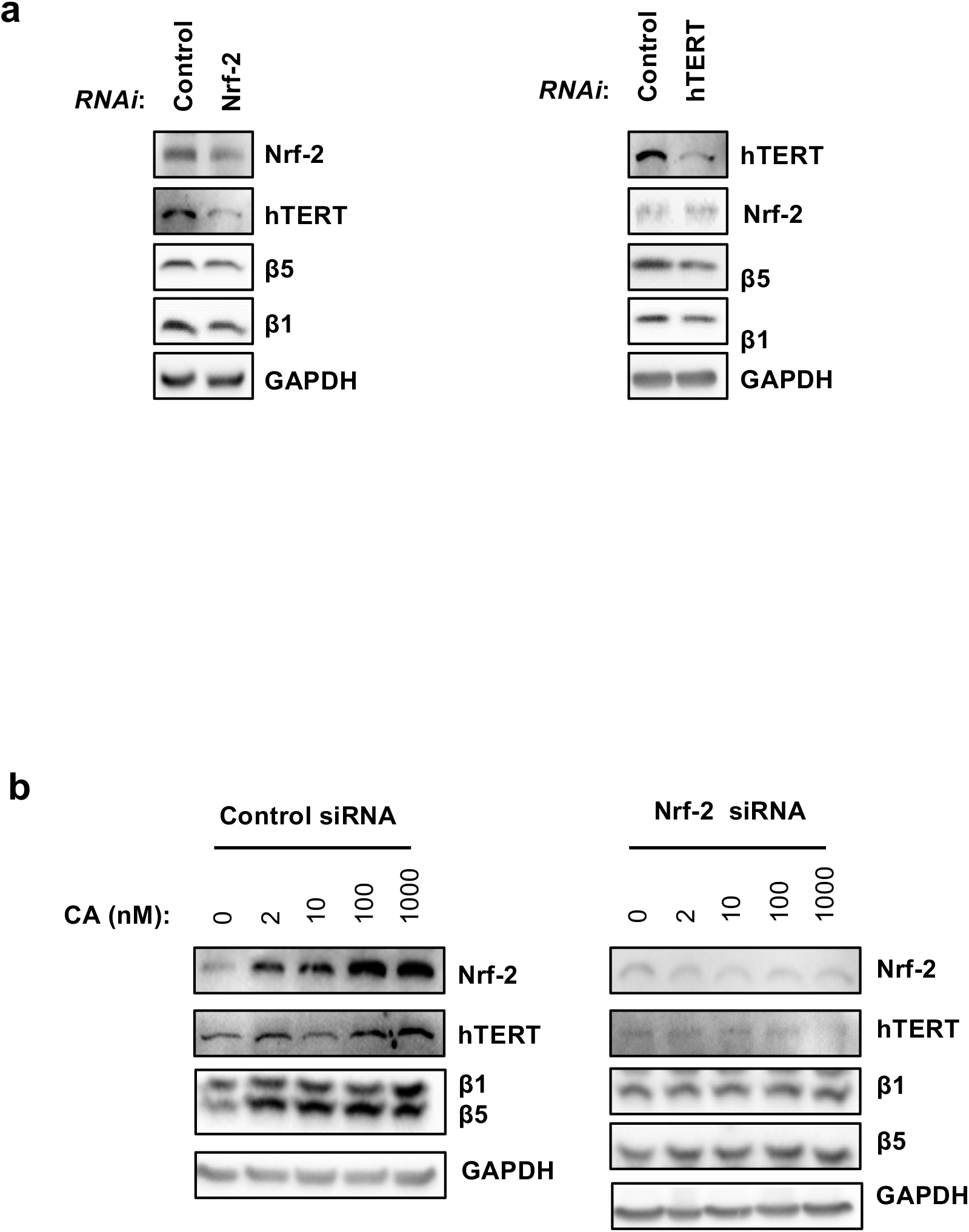
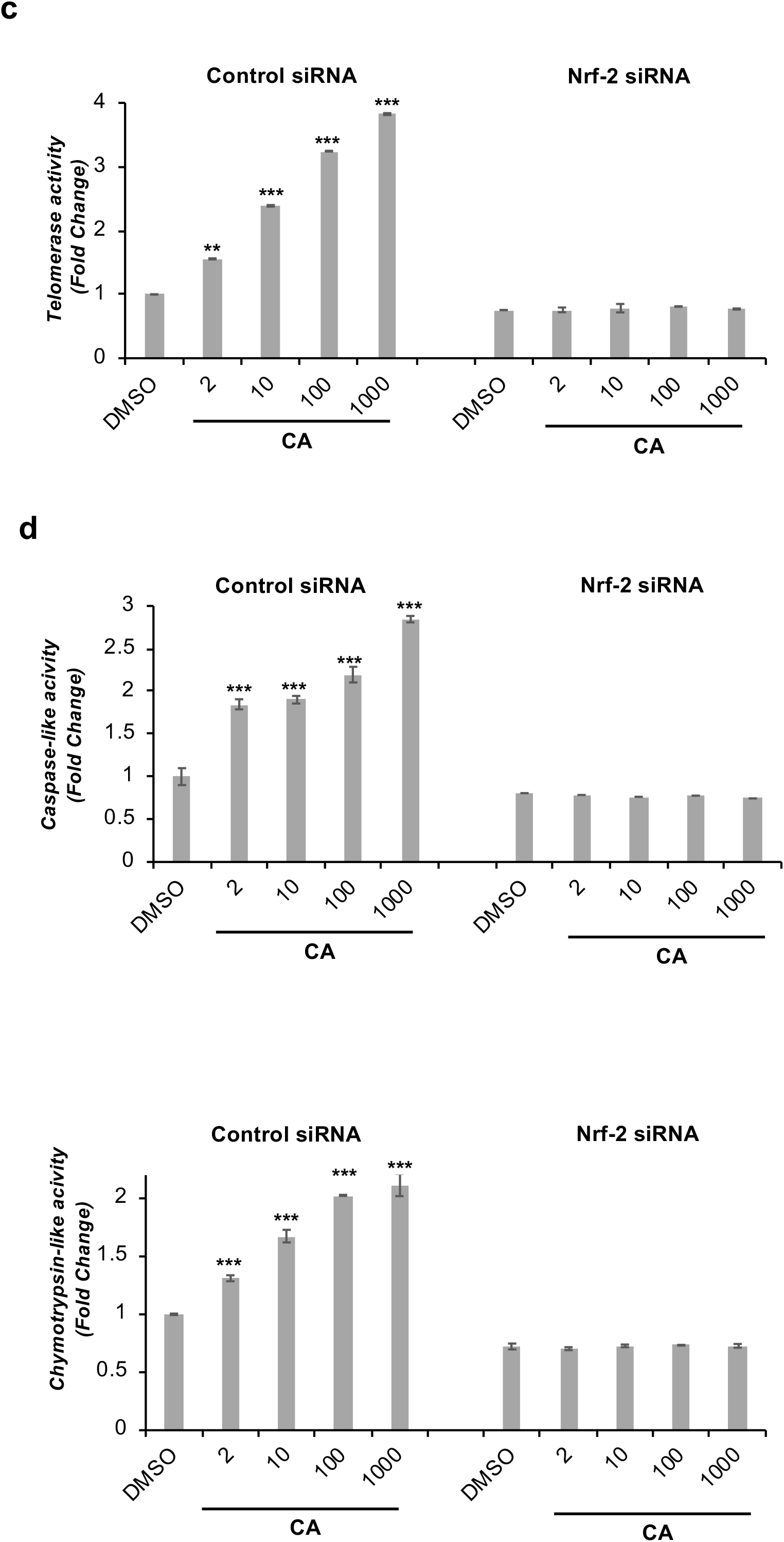
The proteasomal activation of CA is mediated by Nrf-2. **(a)** HEKn cells were transfected with either oligo duplex specific to Nrf-2, hTERT, or Control siRNA. After 48 h later, cells were harvested, and total protein levels of hTERT, Nrf-2, β1, and β5 were determined by IB. GAPDH was used as a loading control. (**b, c, d**) HEKn cells were transfected with either Nrf-2 or Control siRNA (each of them 2 nM). After 48 h later, cells were treated with indicated concentrations of CA for 24 h. (**b**) The total protein levels of Nrf-2, hTERT, β1, and β5 were determined by IB. (**c**) The telomerase enzyme activity was determined by using ELISA (n:3, *p ≤ 0.05, **p ≤ 0.001, ***p ≤ 0.005). (**d**) The proteasomal β1 and β5 subunit activities were determined by using fluorogenic substrates. The activity data were normalized to cellular total protein level and presented as a fold change compared to control cells treated with DMSO used as a solvent control (n:3, *p ≤ 0.05, **p. ≤ 0.001, ***p ≤ 0.005).

**FIGURE 5.**
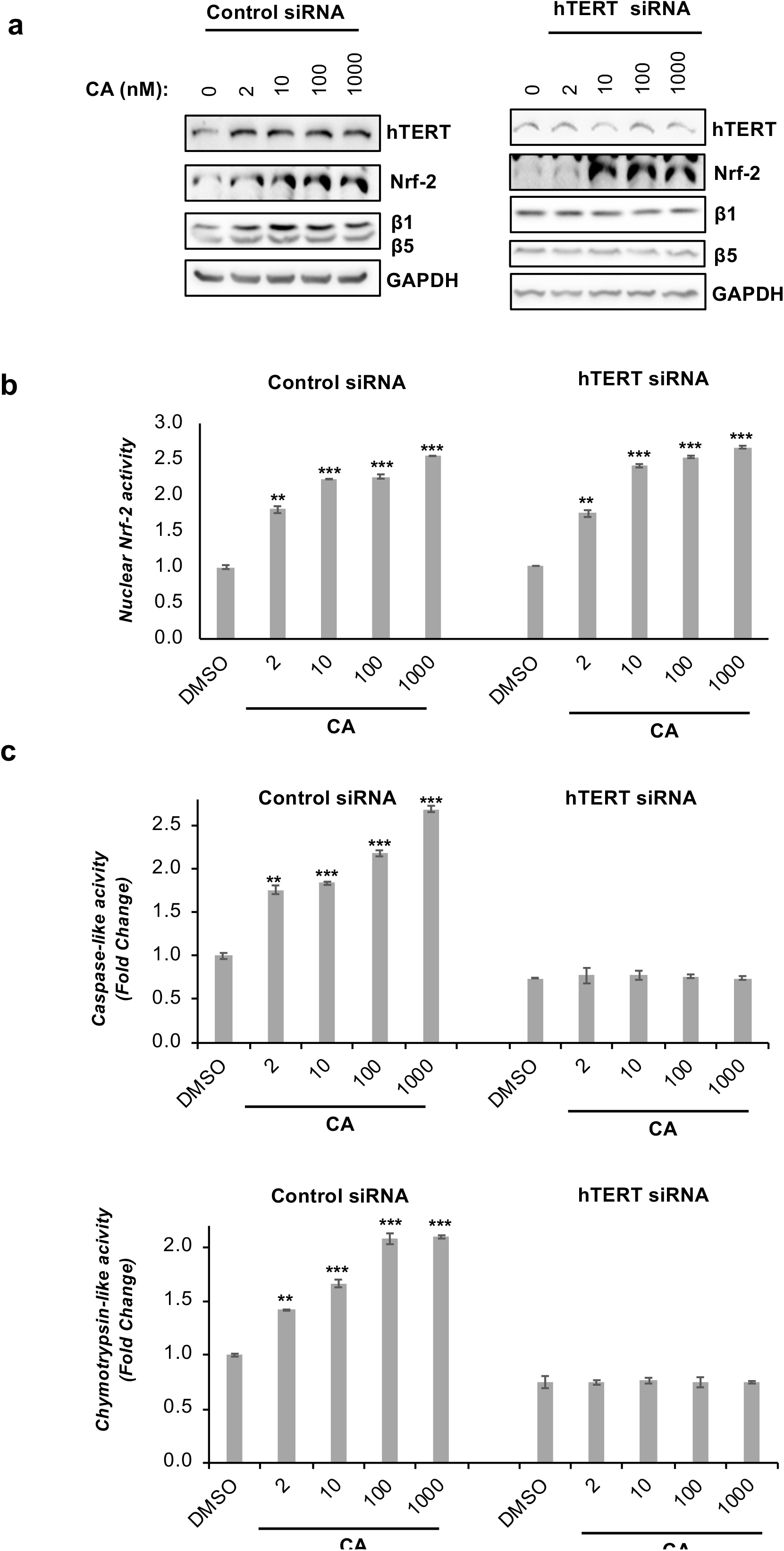
The proteasomal activation of CA is also mediated by hTERT. HEKn cells were transfected with either oligo duplex specific to hTERT or Control siRNA. After 48 h later, cells were treated with CA for 24 h. (**a**) The total protein levels of Nrf-2, hTERT, β1, and β5 were determined by IB. (**b**) Before performing the nuclear Nrf-2 activity, cellular fractionation was performed. The Nrf-2 activity was determined by ELISA. The Nuclear activity data was normalized to nuclear protein level and presented as a fold change compared to control cells treated with DMSO used as solvent control. Error bars are presented as standard deviations (n = 3; *p ≤ 0.05, **p ≤ 0.001, ***p ≤ 0.005). (**c**) The proteasomal β1 and β5 subunit activities were evaluated *via* fluorogenic substrates. The data was normalized to cellular total protein level and presented as a fold change compared to control cells treated with DMSO used as solvent control. Error bars are presented as standard deviations (n = 3; *p ≤ 0.05, **p ≤ 0.001, ***p ≤ 0.005).

### 2.5. CA attenuates the 6-OHDA induced toxicity *via* Nrf-2 and hTERT

In the literature, there are several studies describing CA as a neuroprotective compound against several neurotoxic agents such as Aβ1-42 peptide (Ikram et al., 2021) and 6-OHDA (6-hydroxydopamine) (Nesil et al., 2011). As our data clearly revealed that CA increased Nrf-2, telomerase, and proteasome activity, which reportedly alleviate neurotoxicity, CA-mediated protection against 6-OHDA was next investigated in the context of the involvement of Nrf-2 and hTERT. To this end, control, Nrf-2, or hTERT-silenced SH-SY5Y cells were pre-treated with CA for 8 hours followed by 6-OHDA treatment for an additional 16 hours. We observed that while CA attenuates the 6-OHDA induced toxicity in SHSY5Y cells, its dose-dependent protective effect was abolished in Nrf-2 or hTERT silenced cells (**Figure 6** and **Figure S6**). These results suggest that both Nrf-2 and hTERT activation are essential for CA-dependent protection against 6-OHDA toxicity.

**FIGURE 6.**
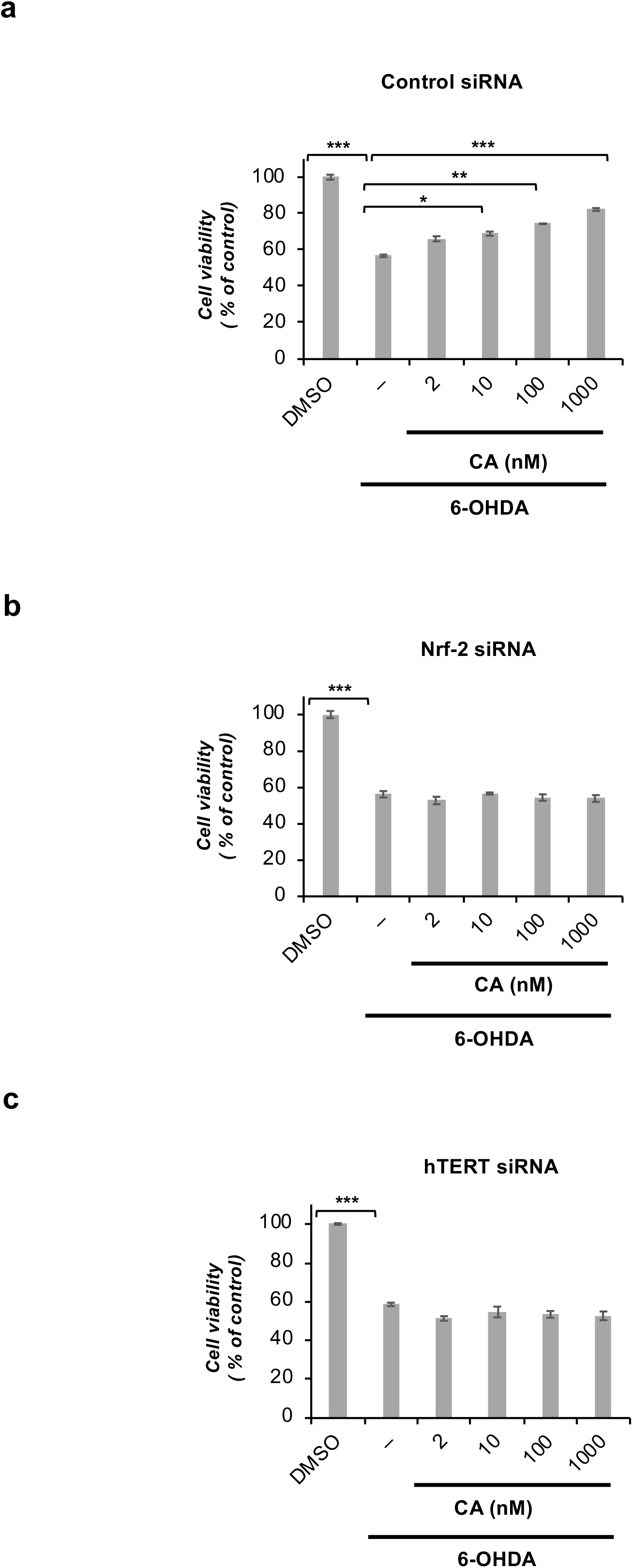
The neuroprotection of CA against 6-OHDA is mediated by both Nrf-2 and hTERT. (**a, b, c**) HEKn cells were transfected with Nrf-2, hTERT, and Control siRNA. After 48 h later, cells were pre-treated with the desired concentration of CA for 8 h, then treated with 6-OHDA for 16 h. The cell viability was assessed by MTT reagent. The absorbance value of cells treated with DMSO as a solvent control for each experiment was considered 100%, and the viabilities of others were calculated. This assay was performed by triplicate samples. Student’s t-test was used to determine the significance of the differences Error bars are presented as standard deviations (n = 3; *p ≤ 0.05, **p ≤ 0.001, ***p ≤ 0.005).

## 3. DISCUSSION

*Radix Astragali* extract was found to have potential to improve population doubling time of human fetal lung diploid fibroblast and fetal kidney cells *in vitro* in the 1970s (Wang, P.C. et al., 2008), and ingredients of the extract have been fully investigated to determine the compounds responsible for delaying senescence and having anti-aging effects as well as wound healing, and anti-inflammatory properties (de Jesus et al., 2011; Sevimli-Gur et al., 2011; Yesilada et al., 2005). One of the major active compounds that are extracted from *Radix Astragalus membranaceus* is Astragaloside IV (AS-IV, 3-O-beta-D-xylopyranosyl,6-O-beta-D-glucopyranosyl-cycloastragenol). In the literature, there are several studies showing that AS-IV protects cells against ROS (Li, H. et al., 2018), ischemia/reperfusion damages (Gu, D.M. et al., 2015), lung injury (Zhang, Z. et al., 2018), and cardiac hypertrophy (Nie et al., 2019) *via* Nrf-2 activation. The other major compound, CA, which is an aglycone and *in vivo* metabolite of AS-IV was discovered as a telomerase activator by TRAP (Telomere Repeat Amplification Protocol) assay (Ip et al., 2014). Furthermore, CA has been implicated in the regulation of various cellular processes as comforting ER stress-related to inflammasome activation (Zhao et al., 2015), improving autophagy *via* inhibition of AKT1-RPS6KB1 signaling (Wang, J. et al., 2018), alleviating hepatic steatosis by activation of farnesoid X receptor signaling (Gu, M. et al., 2017), activation of Src/MEK/ERK pathway (Yung et al., 2012) and upregulation of SIRT1 to prevent neuroinflammation in the ischemic brain (Li, M. et al., 2020). Importantly, CA not only increases telomerase activity and elongates telomere length but also extends life span of mice without augmenting cancer incidences (Bernardes de Jesus et al., 2011). Furthermore, in clinical studies, CA (active ingredient of TA-65®) treatment showed no adverse effects but reduced blood sugar, insulin, total cholesterol, body mass index, and blood pressure (Harley et al., 2013). The decrease in the ratio of total antioxidant capacity and 8-isoprostane levels, which is a hallmark for oxidative lipid damage, suggests that TA-65® may affect redox homeostasis.

All previous reports show that CA is a promising agent in the treatment of age-related diseases, but the molecular mechanisms for how CA influences aging-related main cellular processes such as telomerase activity, redox balance, and proteostasis have not been fully established. Previously, CA was suggested to ameliorate the decrease in Nrf-2 expression caused by TNF-α treatment in vascular smooth muscle cells. In another study, 50 μM CA slightly increased the Nrf-2 transcription in HepG2 cancer cells (Feng et al., 2015), which has much higher intrinsic telomerase expression compared to normal human cells (Sakaguchi et al., 2017; Watanabe et al., 2011). In this study, we found that CA not only increased Nrf-2 protein expression but also enhanced the nuclear localization of Nrf-2, leading to ARE-promotor activity and consequently increasing the levels of cytoprotective enzymes such as HO-1, GR, and GCLC to protect cells against oxidative stress (**Figure 1**).

CA is well-known as a telomerase activator, but its effect on hTERT levels and subcellular localization, as well as their underlying mechanisms, have not been elucidated. Here, in addition to potentiating hTERT expression, we have substantiated that CA increases the nuclear transport of hTERT (**Figure 2c**). Several regulatory proteins play essential roles in the process of hTERT transport to the nucleus. The molecular chaperons p23 and Hsp90 are required to maintain an appropriate hTERT conformation that enables its nuclear translocation (Forsythe et al., 2001; Holt et al., 1999). Moreover, the transport of hTERT along microtubules to the nucleus is facilitated by FKBP52, which links the hTERT-Hsp90 complex to the dynein-dynactin motor (Jeong, Y.Y. et al., 2016). On the other hand, CHIP physically associates with hTERT in the cytosol and prevents its nuclear translocation by disassociating p23, suggesting that CHIP can remodel the hTERT-chaperone complexes. The resulting cytoplasmic hTERT is ubiquitinated by CHIP and subsequently degraded by the proteasome (Lee et al., 2010). Our results revealed that CA enhanced the levels of proteins that function in the nuclear localization of hTERT while decreasing the expression of CHIP, which is known to prevent hTERT nuclear localization and increase hTERT degradation (**Figure 2d,e**). Thus, our data suggest that CA not only increases the expression of hTERT but also enhances its nuclear localization through increasing the levels of Hsp90-based telomerase-associated chaperone complex and augmenting the interaction of hTERT with this complex to contribute to the telomerase activity (**Figure 2 a-f**). The nuclear localization of hTERT is a critical process for the formation of catalytically active telomerase that functions on telomere maintenance. Interestingly, oxidative stress induces nuclear export of hTERT (Ahmed et al., 2008; Haendeler et al., 2003). Moreover, upon H_2_O_2_ treatment, no degradation of hTERT protein was observed, but rather a concomitant increase within the mitochondria (Ahmed et al., 2008; Haendeler et al., 2003). Supporting this view, we also observed that the mitochondrial hTERT levels increased upon H_2_O_2_ treatment (**Figure 2g**). hTERT was found to be associated with mitochondrial DNA (mtDNA) as well as mitochondrial tRNA and localized in the mitochondrial matrix (Haendeler et al., 2009; Sharma et al., 2012), where it should have non-telomeric functions as circular mtDNA does not contain telomeres. The function of hTERT in mitochondria has been controversially reported (Rosen et al., 2020). Aggravated mtDNA damage and apoptosis were reported in TERT-overexpressing fibroblast cell lines upon oxidative stress (Santos et al., 2006). Conversely, it has also been demonstrated that TERT protects mtDNA and reduces mitochondrial ROS production suggesting a protective role for TERT in mitochondria (Haendeler et al., 2009). When examining the effect of CA on the subcellular localization of hTERT in the presence of oxidative stress, we observed that CA attenuated the H_2_O_2_-induced mitochondrial hTERT levels. Furthermore, the nuclear hTERT level was increased with CA treatment compared to solvent control, even in the presence of oxidative stress (**Figure 2g**). Considering the effect of CA to alleviate the H_2_O_2_-induced ROS production and its neuroprotective activities, it can be speculated that the reduction of mitochondrial hTERT levels by CA might be due to its cellular protective roles *via* Nrf-2/ARE against oxidative stress.

The proteasome genes are among the known-downstream targets of Nrf-2 pathway (Kwak et al., 2003), and Nrf-2 increases proteasome subunit protein expression and proteasome activity (Arlt et al., 2009; Kapeta et al., 2010). Moreover, hTERT was determined to bind to multiple proteasome subunits and promote 26S proteasome assembly, which indicates another non-canonical function for hTERT that is independent of its telomere-regulatory function (Im et al., 2017). Studies toward the discovery of proteasome activators have been reported several natural and synthetic small compounds that stimulate proteasomal activity (Chondrogianni et al., 2020). Although there are several studies regarding proteasome activation of triterpenoids such as Betulinic acid (Huang et al., 2007), 18α-GA (Kapeta et al., 2010), and Hederagenin (Kapeta et al., 2010), no study has been reported to determine the effects of CA on the proteasomal activity. As the regulation of proteasomal activity and/or assembly was found to be associated with both telomerase, and Nrf-2 (Imai et al., 2003; Kapeta et al., 2010), the effect of CA on proteostasis was investigated. Here, we presented that CA not only enhances the activity of β1 and β5 proteasome subunits, responsible for caspase-like and chymotrypsin-like catalytic activities, respectively but also increases the mRNA/protein levels of these subunits and assembly of proteasome (**Figure 3b-e**). With the results obtained from the present study, we propose that CA, known as a small molecule telomerase activator, would also be considered as a potent activator of Nrf-2 and proteasome.

In this study, we report for the first time that CA treatment affects multiple aging-related cellular processes such as hTERT, Nrf-2, and proteasome. In addition to the interrelationships of proteasome-hTERT and proteasome-Nrf-2 system, there are several studies about the bilateral relations between hTERT and Nrf-2. XRCC5 (X-ray repair cross-complementing 5), a transcriptional regulator factor binding to the hTERT promoter, was reported to physically interact with Nrf-2, which in turn upregulated hTERT in an Nrf-2 dependent manner in hepatocellular carcinoma (HCC) cells (Liu et al., 2022). Consistently, administration of DMF, an activator of Nrf-2 pathway, to mice resulted in significant elevation of TERT mRNA levels (Akino et al., 2019). TERT was also found significantly reduced in lung tissue of Nrf2^-/-^ mice in the model of intestinal ischemia/reperfusion-induced acute lung injury (Dong et al., 2021). On the other hand, hTERT G4-targeting small molecule (RG1603), or pharmacological inhibitor of hTERT (costunolide) and siRNA-mediated depletion of hTERT expression resulted in inhibition of Nrf-2 and HO-1 expression in glioblastoma and prostate cancer (Ahmad et al., 2016; Song et al., 2019). Interestingly, hTERT was found to be primarily upregulated Nrf-2 by increasing Nrf-2 promoter activity rather than by regulating Nrf-2 mRNA or protein stability in colon cancer cells (Gong et al., 2021). Considering our results, which revealed that CA increased both Nrf-2, telomerase, and proteasome activity, we aimed to identify the underlying mechanism in the next step. First, we found that Nrf-2 silencing caused downregulation of hTERT protein levels, but hTERT silencing had no significant effect on Nrf-2 protein levels (**Figure 4a**). Second, we found that CA lost its ability to increase the protein levels and activities of hTERT, β1, and β5 proteasomal subunits in Nrf-2 silenced HEkn cells (**Figure 4b-d**). On the other hand, in hTERT-silenced HEKn cells, CA still retained its ability to increase Nrf-2 activity but lost its effect on the proteasome (**Figure 5**). Considering all these data together, it can be suggested that the telomerase activator effect of CA may be dependent on Nrf-2 activity or due to the significant decrease in hTERT expression observed in Nrf-2 silenced cells, CA cannot be able to increase the telomerase activity in Nrf-2 silenced cells.

CA treatment exhibited stimulatory effect on telomerase activity, which was consistent with augmented TERT expression and nuclear translocation. Additionally, CA robustly augmented expression of ARE signaling molecules, including Nrf-2, HO-1, GR, and GCLC. Thus, CA could antagonize oxidative/electrophilic stress and DNA damage while improving telomere function, which are known to be important in neurodegenerative diseases. On the other hand, neurodegenerative diseases such as Parkinson’s, Alzheimer’s, and Huntington’s have been associated with intracellular ubiquitin-positive inclusions formed by aggregate-prone neurotoxic proteins. It has been reported that the aberrant ubiquitin-proteasome system in neurodegenerative diseases contributes to the accumulation of neurotoxic proteins that perturb cellular homeostasis and neuronal function and to instigate neurodegeneration (Wang, P.C. et al., 2008; Zhang, D.D. & Hannink, 2003; Zheng et al., 2016). Thus, enhancement of proteasome activity by small molecules has been considered as a new approach for treatment of neurodegenerative diseases (Sevimli-Gur et al., 2011). Based on the data observed from this study suggesting that CA potently activates proteasome through activation of hTERT and Nrf-2 systems, we analyzed the molecular mechanism of reported neuroprotection properties (Nesil et al., 2011; Wan et al., 2021) of CA by utilizing 6-OHDA toxicity in Nrf-2 and hTERT silenced HEKn cells. While CA provided neuroprotection against 6-OHDA in a dose-dependent manner, this neuroprotection was abolished in both hTERT, and Nrf-2 silenced HEKn cells suggesting that the neuroprotective effect of CA is driven by these pathways (**Figure 6**).

This study is the first to report that CA has great potential as a proteasome activator in addition to its role as telomerase and Nrf-2 activator to decelerate the progression of aging and age-related diseases. Furthermore, our results provide new evidence for the molecular mechanism by which CA acts at the intersection of three aging-related cellular processes, namely telomerase, Nrf-2, and proteasome.

## 4. EXPERIMENTAL PROCEDURES

### 4.1. Reagents and Antibodies

While DMF (242926), 18α-GA (G8503), 6-OHDA (H4381), and DMSO (Dimethylsulfoxide; D2650) were purchased from Sigma Aldrich, GA (13355) was obtained from Cayman Chemical, and Mg-132 (474790) was obtained from Calbiochem. The proteasome fluorogenic substrates (Suc-LLVY-AMC-BML-P802-0005 for the ChT-L, Z-LLE-AMC-BML-ZW9345-0005 for the C-L and Boc-LRR-AMC-BML-BW8515 for the T-L) and primary proteasome antibodies against β1 (BML-PW8140), β2 (BML-PW8145), β5 (BML-PW8895), α4 (BML-PW8120), and α6 (BML-PW8100) were purchased from Enzo Life Sciences. Antibodies against Nrf-2 (Abcam, Ab-62352), Nrf-2 (Bioss, bsm-52179R), hTERT (Origene, TA301588), Keap-1 (CST-8047), HO-1 (CST-5853), Hsp90 (Santa Cruz, sc-69703), p23 (Santa Cruz, sc-101496), CHIP (Santa Cruz, sc-133066), FKBP52 (Santa Cruz, sc-100758) and p16 (CST, 80772) were used in this study. Antibodies against β-actin (Sigma-Aldrich-A5316), GAPDH (CST, 5174), α-Tubulin (CST, 3873), VDAC (CST, 4661), and Lamin A-C (CST, 4777) were used as loading control. Secondary antibodies (Goat anti-rabbit-31460 and Goat anti-mouse-31430) were purchased from Thermo Fisher Scientific.

### 4.2. Cell line and Culture conditions

HEKn (human neonatal epithelial keratinocyte cells) cells were obtained from the American Type Culture Collection (ATCC; PCS-200-010), and cultured in Dermal Cell Basal Media (ATCC; PCS-200-030) supplemented with Keratinocyte Growth Kit (ATCC; PCS-200-040) according to supplier’s instructions at 37°C under humidified 5% CO_2_. HEKn cells were sub-cultured when they reached 70-80% confluency.

SHSY-5Y (human neuroblastoma) cells were obtained from the American Type Culture Collection (ATCC; CRL-2266), and cultured in DMEM (Dulbecco’s modified Eagle’s medium; Lonza, BE12-604F) supplemented with 10% FBS (fetal bovine serum, Biological Ind., Israel, 04-007-1A) and 2 mM L-glutamine (Biological Ind., Israel, 03-020-1B) at 37°C under humidified 5% CO_2_.

### 4.3. Cellular fractionation

HEKn cells were treated with indicated doses of CA or DMSO for 24 h. Nuclear-cytoplasmic fractionation was performed according to the NE-PER kit (Pierce, Thermo Fisher Scientific, US, 78833). The nuclear lysates were subjected to Nrf-2 Transcription Factor Assay (Cayman Chemical, US, 600590) to evaluate the effect of CA on the nuclear Nrf-2 activity. Also, the cytosolic and nuclear lysates were used for the nuclear translocation of both Nrf-2 and hTERT. In these experiments, GAPDH and α-Tubulin were used as cytoplasmic loading controls and Lamin A-C as a nuclear loading control.

Mitochondrial and nuclear cellular fractions were performed *via* Mitochondrial isolation kit (Thermo Fisher Scientific, US, 89874), as previously reported (Stasik et al., 2008). Briefly, the cells were lysed in lysis buffer, then centrifuged at 800 g for 10 min. After centrifugation and washing steps with 1x PBS, the nuclear pellet was lysed in 1% nonidet P-40, 0.5% sodium deoxycholate, and 0.1% SDS in 1X PBS, pH 8.0 and centrifuged at higher speed as 16000x*g* for 20 min. The supernatant containing mitochondrial and cytoplasmic fractions was centrifuged at higher speed as 12000x*g* for 10 min. The pellet was dissolved in 1% nonidet P-40, 0.5% sodium deoxycholate, and 0.1% SDS in 1X PBS, pH 8.0, and centrifuged to obtain soluble mitochondrial protein. After performing cellular fractionation, nuclear and mitochondrial protein levels were determined by BCA protein assay (Thermo Fisher Scientific, US, 23225). In these experiments, while VDAC was used as mitochondrial loading controls, Lamin A-C was used as a nuclear loading control.

### 4.4. Immunofluorescence Analysis

HEKn cells were seeded on 6 well plate with coverslips for fluorescence microscopy (Uner et al., 2020). The next day, cells were treated with the indicated concentration of CA for 24 h. Then, cells on coverslips were washed twice with cold 1X PBS and fixed with 4% paraformaldehyde in PBS for 30 min at 4°C. After washing six times with PBS, cells were blocked with 0.01% saponin and 0.01% BSA in PBS for cell permeabilization. The samples were first incubated with anti-Nrf-2 antibody (Abcam, Ab-62352, 1:250 dilution) at 4°C overnight and then with Alexa Flour secondary antibody (Pierce, A-11008, 1:500 dilution) for 1 h at 37°C. Slides were mounted via mounting media with DAPI (Invitrogen, P36965) for nuclei staining. Images were taken by fluorescence microscopy (Olympus IX70, Japan).

### 4.5. Luciferase activity assay

HEKn cells were transfected with pGL4.37[luc2P/ARE/Hygro] Vector (Promega, US, E364A) using Turbofect transfection reagent (Thermo Fisher Scientific, US, R0532) for 24 h followed by treatment of indicated concentrations of CA for 24 h. DMF (50 μM) was used as an experimental control. Cells were harvested and lysed in Cell Culture Lysis Reagent (Promega, US, E1500), and supernatants were used for luciferase activity assay according to the manufacturer’s instruction. Standard deviations and means were obtained from three independent biological replicates.

### 4.6. Cytoprotective enzyme activity assay

HEKn cells were seeded at 60 mm culture dish. The next day, cells were treated with the indicated concentration of CA and DMF (50 μM), which is used as an experimental control. for 24 h. HO-1 (ab207621), GCLC (ab233632), and GR (Cayman Chemical, 703202) enzyme activities were determined according to the manufacturers’ instruction.

### 4.7. Proteasome activity assays

ChT-L, C-L, and T-L activities were assayed in crude extracts incubated with CA for 1 h or lysates obtained from cells were incubated with CA for 24 h under culture conditions as described previously. To determine proteasome subunit activities of CA, cells were lysed (with a buffer containing 8,56 g sucrose, 0.6 g HEPES, 0.2 g MgCl_2_, 0.037 g EDTA in ml; DTT (dithiothreitol) was added freshly at a final concentration of 1 mM (Karademir et al., 2018). The supernatant was incubated with reaction mixture containing 7.5 mM MgOAc, 7.5 mM, MgCl_2_, 45 mM KCl, and 1 mM DTT for 10 min (Karademir et al., 2018) and then with fluorogenic substrates Suc-LLVY-AMC (Enzo Life Sciences, BML-P802-0005), Z-LLE-AMC (Enzo Life Sciences, BML-ZW9345-0050), and Boc-LRR-AMC (Enzo Life Sciences, BML-BW8515-0005) for 1 h to determine proteasome subunit activities. The measurements were performed by a fluorescent reader at 360 nm excitation/460 nm emission (Varioscan, Thermo Fisher Scientific, US). While DMSO was used as solvent control, Mg-132, known as a proteasome inhibitor, was used as an experimental control. Protein concentrations were determined using the BCA (bicinchoninic acid) assay (Thermo Fisher Scientific, USA).

### 4.8. Measurement of Reactive Oxygen Species

To detect ROS formation, 2’-7’-Dichlorodihydrofluorescein diacetate (H_2_DCFDA) (Enzo; ALX-610-022-M050) was used as previously reported (Gezer et al., 2022). For this purpose, HEKn cells were pre-treated with indicated concentrations of CA for 1 h. After 1 h pre-treatment of CA, cells were treated with 250 μM H_2_O_2_ (hydrogen peroxide) for 24 h. H_2_DCFDA resuspended in 1X PBS at a final concentration of 10 μM was added to the cells and incubated at 37 °C for 30 min (Kapeta et al., 2010). After the incubation period, half of the cells were suspended, and execution/emission absorption of the oxidative product was measured at 495 and 520 nm. The other half of cell suspension was used to determine protein concentration by BCA assay to perform normalization of absorbance. This experiment was performed in three independent experiments.

### 4.9. Immunoblotting (IB)

Cell lysates were prepared in RIPA buffer (1% nonidet P-40, 0.5% sodium deoxycholate, and 0.1% SDS in 1X PBS, pH 8.0). with protease inhibitors (Roche, Switzerland). Protein concentrations were determined by BCA assay (Thermo Fisher Scientific, US). Equal amounts of samples were loaded to the gels and separated by SDS-PAGE electrophoresis after 5-min treatment with 4X Laemmli buffer at 95 degrees. Then, gels were transferred to the PVDF membrane (EMD Millipore, US, IPVH00010). Membranes were blocked with 5% non-fat dry milk prepared in 1X PBS-0.1% Tween-20, then incubated with primary and secondary antibodies. β-actin and GAPDH were used as loading control for whole lysate experiments. Chemiluminescence signals of the proteins were detected with Clarity ECL substrate solution (BIORAD, US, 1705061) by Vilber Loumart FX-7 (Thermo Fisher Scientific, US). The quantifications and analysis of proteins were determined by ImageJ software (http://imagej.nih.gov/ij/). All IB experiments were performed at least in three independent replicates.

### 4.10. Real-Time PCR Analysis

RNA isolations were performed by using Aurum Total RNA Mini Kit (Bio-Rad, USA, 7326820) according to the supplier’s instructions. RNA concentrations were measured by using Beckman Coulter Du730 instrument. 1 μg RNA of each sample was used for cDNA synthesis. cDNA synthesis was performed *via* Iscript cDNA Synthesis Kit (Bio-Rad, USA, 1708891). Gene expression analysis was performed by using SYBR Green 1 (Bio-Rad, USA, 1725121) and LightCycler480 Thermocycler (Roche, Switzerland). For RT-PCR studies, specific primers against Nrf-2, hTERT, β1, and β5 (**Table S1**). While 300 nM primer pairs of Nrf-2, β1, and β5 were used in 10 μl reactions, 500 nM primer pairs were used for hTERT. Fold changes of the transcripts were done by the normalizations against housekeeping gene as GAPDH. For the analysis of Ct values, Qiagen Rest 2009 program was used. Gene expression experiments were performed at three independent biological replicates with three technical replicates.

### 4.11. RNA Interference

Specific siRNA oligo duplexes and Control siRNA (Origene, USA, SR30004) were complexed with Lipofectamine 2000 (Invitrogen, USA, 11668019) and serum-free Opti-MEM (Invitrogen, USA, 31985070) according to the manufacturer’s instruction. When HEKn cells reached at confluency of 60-70%, cells were transfected with 2 and 15 nM siRNA against Nrf-2 (Origene, USA, SR321100) and hTERT (Origene, USA, SR322002), respectively. Equal molar control siRNA was used as experimental control. After 24 h later, media was refreshed and further incubation in complete media for HEKn cells was followed for an additional 24 h. Lastly, cells transfected with Nrf-2, hTERT, or control siRNA were treated with indicated doses of CA for an additional 24 h. In all silencing experiments, to ensure functional silencing of Nrf-2 and hTERT, the controlling of these proteins’ levels was revealed by IB.

### 4.12. Immunoprecipitation

HEKn cells were treated with desired concentrations of CA for 24 h. Then, cells were harvested and lysed in IP buffer containing 50 mM Tris.Cl pH:8, 150 mM NaCl, 1 mM EDTA, 0.2 % NP-40. Protein amounts were performed by BCA assay. 1-2 μg anti-α6 or anti-Hsp90 antibody was added to lysates with equal protein concentrations and incubated on a rotator at 4 °C overnight. Protein A-agarose beads (Invitrogen, USA, 1041) that were pre-washed with lysis buffer were added to lysate-antibody complexes and incubated on a rotor at 4 °C overnight. Immunoprecipitated proteins were collected by centrifugation and washed five times with Lysis Buffer and eluted from beads by boiling for 5 min in 1X Laemmlli Buffer. After immunoprecipitation, the samples were processed for IB.

### 4.13. Telomerase activity assay

The determination of telomerase enzyme activity was done in HEKn cells by using TeloTAGGG Telomerase PCR Plus Kit (Sigma Aldrich, USA, 12013789001) according to the manufacturer’s instruction. Values are presented as fold change (relative value of DMSO used as a solvent control). Experiments were performed at least in three independent biological replicates.

### 4.14. Cell viability assay

The neuroprotective effect of CA against 6-OHDA was evaluated by MTT reagent (3-(4,5-Dimethyl-2-thiazolyl)-2,5-diphenyl-2H-tetrazolium bromide) (Sigma Aldrich, UK, M5655) according to the manufacturer’s instruction. For this purpose, SHSY-5Y cells were transfected with hTERT and Nrf-2 specific oligo duplex (2 nM for Nrf-2, 15 nM for hTERT duplex) or control siRNA. After 48 h, cells were pre-treated with indicated doses of CA for 8 h. Then, cells were treated with 6-OHDA for another 16 h (Hu et al., 2014). At the end of the incubations, the mixture of MTT and medium (1:9) was replaced with old media. The formazan crystals were dissolved in DMSO. The absorbance was measured with a microplate reader at 570 nm (Varioscan, Thermo Fisher Scientific, US) and 690 nm as a reference wavelength. Cell viability was shown as a percentage cell viability compared to cells treated with DMSO as solvent control. The MTT experiment was performed by three different replicates.

### 4.15. Statistical Analysis

Data are exhibited as means± standard deviation (SD). Student t-Test or One-way ANOVA Post Hoc test was used to analyze statistical by using GraphPad Prism software. The significance of the differences was determined as *p ≤ 0.05, **p ≤ 0.001, ***p ≤ 0.005.

## ACKNOWLEDGEMENTS

This study was supported by Scientific and Technological Research Council of Turkey (TUBITAK, Grant number: 119Z086).

We thank the Pharmaceutical Sciences Research Centre (FABAL, Ege University, Faculty of Pharmacy) for equipment support and Bionorm Natural Products for donating Cycloastragenol.

## AUTHOR CONTRIBUTIONS

P.B.K and E.B. hypothesed and designed the study. S.Y. participated in the design of the experiments, conducted the experiments and collected the data. P.B.K, E.B and S.Y. analyzed the results and contributed to the manuscript preparation. All authors read and approved the manuscript.

## ABBREVIATIONS

Nrf-2: nuclear factor erythroid 2-related factor 2
Keap-1: Kelch-like ECH-associated protein 1
ARE: Antioxidant Response Elements
TERT: telomerase reverse transcriptase
TERC: Telomerase RNA Component
CA: Cycloastregenol
DMF: Dimethyl fumarate
18α-GA: 18α-Glycyrrhetinic acid
6-OHDA: 6-hydroxydopamine
GA: geldanamycin
DTT: dithiothreitol
H2DCFDA: 2’-7’-Dichlorodihydrofluorescein diacetate
ROS: reactive oxygen species
H_2_O_2_: hydrogen peroxide
HO-1: heme oxygenase 1
GR: glutathione reductase
GCLC: Glutamatecysteine ligase catalytic subunit
CHIP: C terminus of Hsc70-interacting protein
Hsp90: Heat shock protein 90
FKBP52: FK506-binding protein 52
HEKn: neonatal human primary epidermal keratinocytes
IB: immunoblotting
IP: immunoprecipitation

## SUPPLEMANTARY FIGURE LEGENDS

**Figure S1 (A)** IB analysis of p16 (senescence marker) protein levels in senescent (p15) and young (p5) HEKn cells. (**B**) The effect of CA on nuclear Nrf-2 activity in senescent HEKn. HEKn cells were treated with indicated concertation of CA for 24 h. Nrf-2 transcription factor assay using the nuclear proteins obtained from control or CA-treated senescent HEKn cells were performed by an ELISA kit. The data were normalized to nuclear protein levels and presented as a fold change *vs* control cell (treated with DMSO). Error bars are presented as standard deviations (n = 3; *p ≤ 0.05, **p ≤ 0.001, ***p ≤ 0.005).

**Figure S2.** CA increased hTERT protein levels in senescent HEKn cells. HEKn cells were treated with indicated concertation of CA for 24 h. The protein levels of hTERT were evaluated by IB.

**Figure S3. (3A).** CA decreased CHIP protein levels in senescent HEKn cells. HEKn cells were treated with indicated concertation of CA for 24 h. The protein levels of CHIP were evaluated by IB. (**3B**) CA increases Hsp90-chaperon complex protein levels in MRC-5 cells. The protein levels of Hsp90, p23, FKBP52, and hTERT were determined by IB.

**Figure S4.** The effect of Mg-132 on proteasome-enriched lysate of HEKn cells. Proteasome-enriched lysate of HEKn cells was incubated with 5 μM concentration of Mg-132. The proteasome subunits’ activities were determined *via* fluorogenic substrates. Error bars are presented as standard deviations (n = 3; *p ≤ 0.05, **p ≤ 0.001, ***p ≤ 0.005).

**Figure S5.** The effect of CA on proteasome subunit activities and protein levels in senescent HEKn cells. HEKn cells were treated with the indicated concentration of CA. **(A)** The proteasome subunits’ activities were determined *via* fluorogenic substrates. Error bars are presented as standard deviations (n = 3; *p ≤ 0.05, **p ≤ 0.001, ***p ≤ 0.005). **(B)** The protein levels of β1 and β5 were evaluated by IB.

**Figure S6.** Determination of siRNA transfection efficiency of Nrf-2 and hTERT in SHSY-5Y cells. SHSY-5Y cells were transfected with specific oligo duplex to Nrf-2, hTERT, or Control siRNA. After 72 h later, the cells were harvested, and protein levels of Nrf-2, hTERT, β1, and β5 were evaluated by IB.

**Table S1.** q-RT-PCR primer list

