## supplemental file for "The role of Cycloastragenol at the intersection of Nrf-2/ARE, telomerase, and proteasome activity"

#### Figure S1

A

B

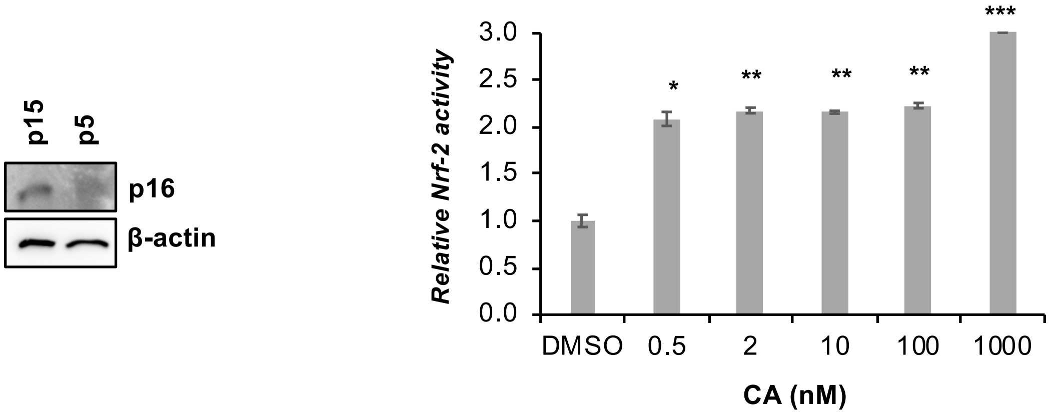


#### Figure S2

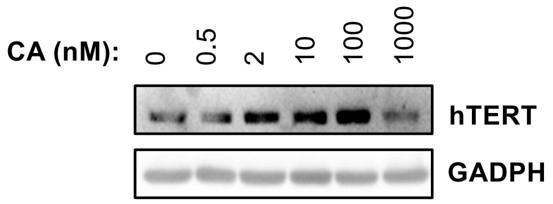


#### Figure S3

A

B

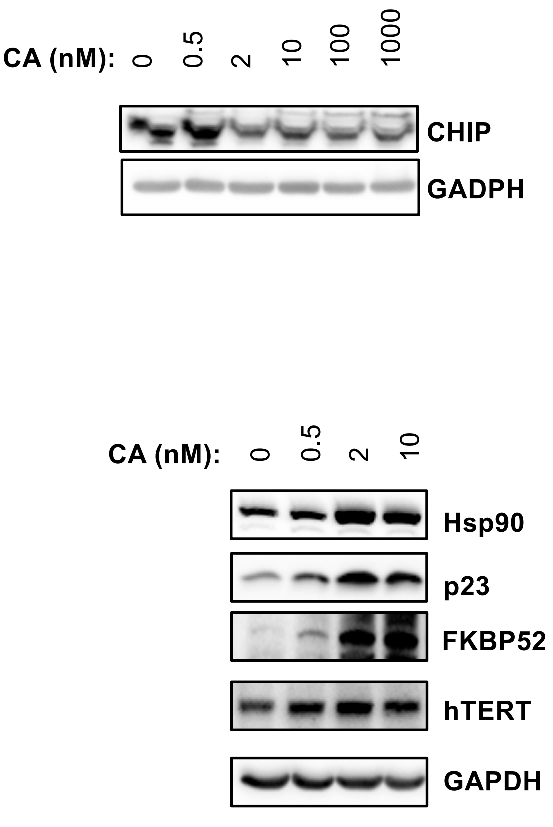

### Figure S4

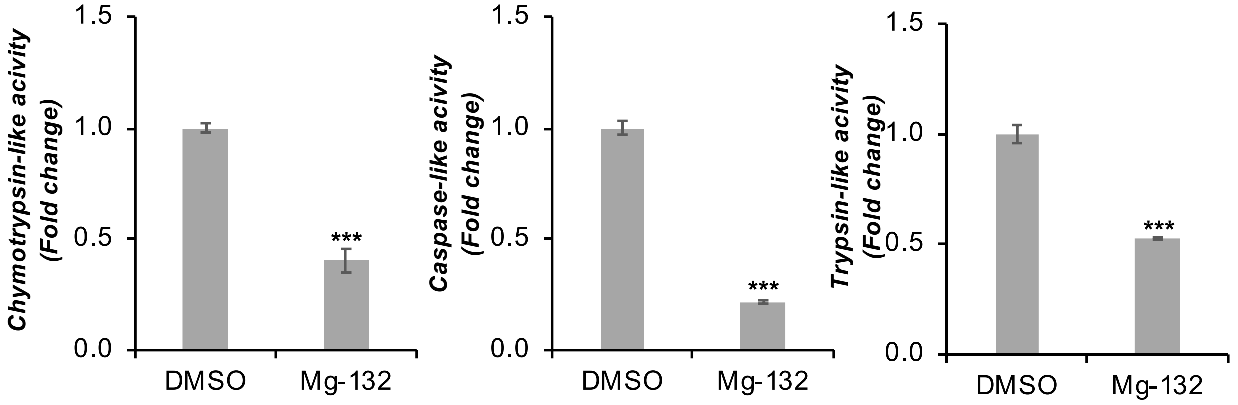


### Figure S5

**A**

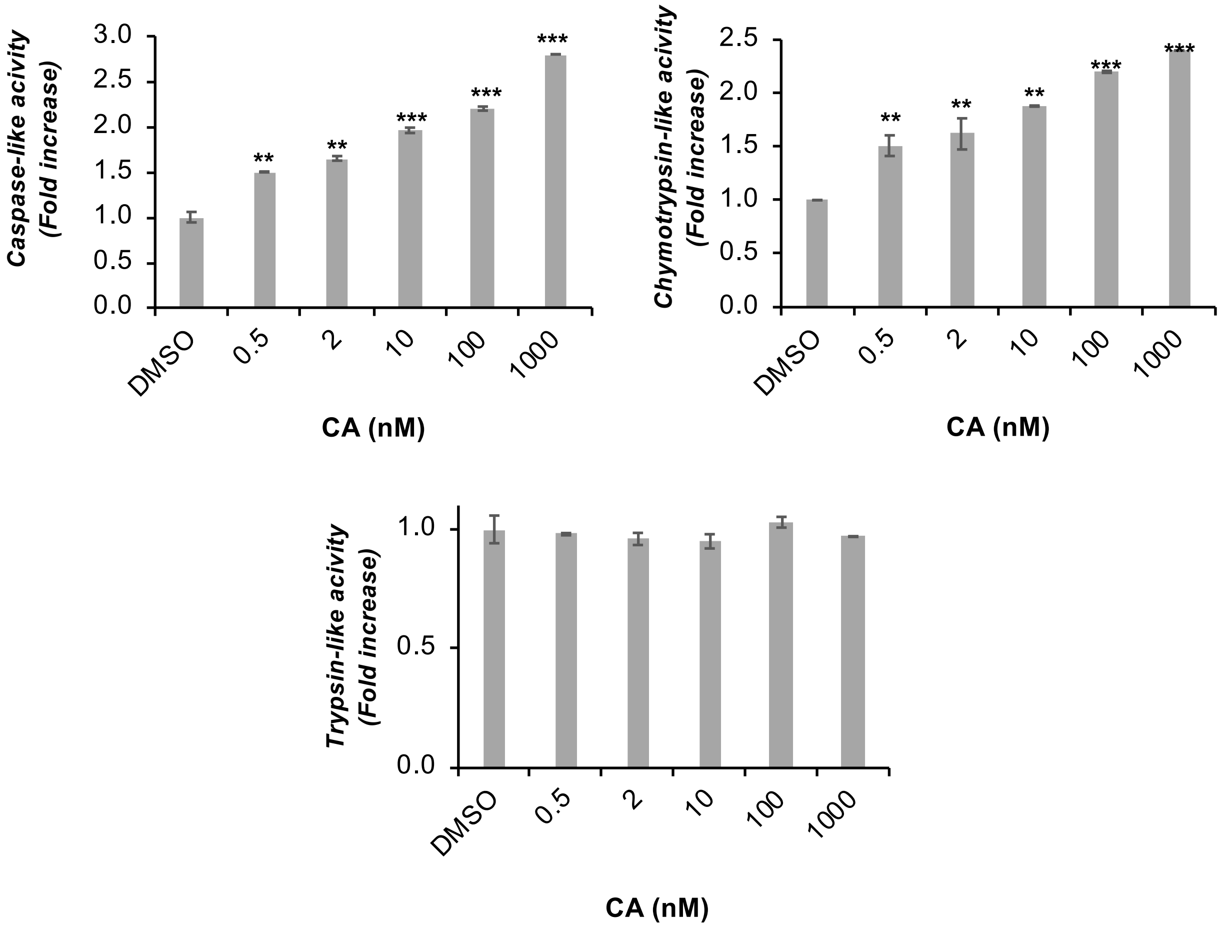


**B**

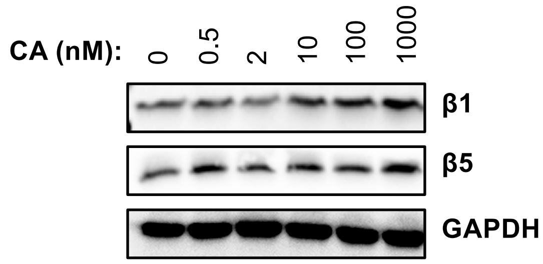

#### Figure S6

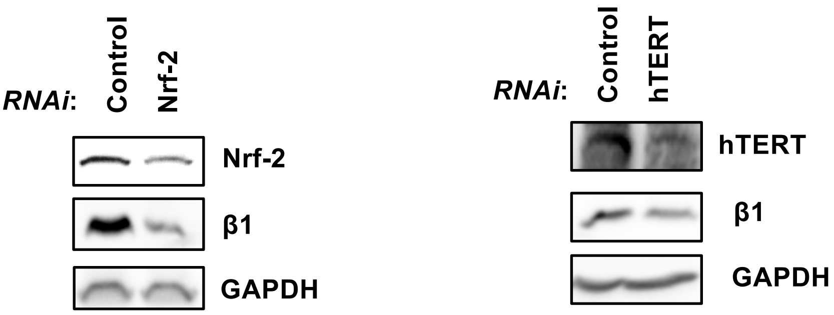


#### Table S1

| Target gene | Forward sequences | Reverse sequences |
|---|---|---|
| Nrf-2 | GTGAATTTCTCCCAATTCAGCCAG | TCAGGAACAAGTGACTGAAACGTA |
| hTERT | CGGAAGAGTGTCTGGAGCAA | GGATGAAGCGGAGTCTGGA |
| β1 | AGACTGTCTTACGCTGACAAAGAT | CATAGTATGGAAAGAAGCGCCTTG |
| β5 | GATCTAGGTCCAGGGAGTCTCAG | GAAGCTCATAGATTCGACATTGCC |
